# Active Nuclear Shuttling Enables Efficient Virus-Free CAR Gene Integration Using Ready-to-Use Lipid Nanoparticles

**DOI:** 10.64898/2026.05.29.728914

**Authors:** Kota Inoue, Misaki Masui, Sayako Umetani, Kazuhiko Nakata, Kohei Shimizu, Kohei Yasuda, Kentaro Numajiri, Yuta Murakami, Shinichi Hashimoto, Hirofumi Fukunaga, Nao Suzuki-Yamazaki

**Author notes:** Corresponding author: Nao Suzuki-Yamazaki, PhD, Bio Science & Engineering Laboratories, FUJIFILM Corporation, Kanagawa, Japan, 577 Ushijima, Kaisei-machi, Ashigarakami-gun, Kanagawa, Japan.

## Abstract

Non-viral engineering of chimeric antigen receptor T (CAR-T) cells is highly desirable to overcome the cost, safety, and scalability limitations associated with viral vectors and electroporation. However, efficient nuclear delivery and stable genomic integration of DNA in primary human T cells remain major challenges. Here, we established a virus-free CAR-T manufacturing platform using lipid nanoparticles (LNPs) combined with a nuclear localization signal (NLS) shuttle strategy. We developed proprietary ready-to-use LNPs that enable on-demand encapsulation of nucleic acids. To overcome nuclear transport barriers, NLS–fused transposase or genome-editing nuclease was used to bind donor DNA in the cytoplasm and promote active nuclear import. This approach enabled highly efficient and low-toxicity delivery of mRNA and plasmid DNA into primary human T cells. NLS–assisted transposase delivery markedly enhanced genomic integration of the CAR gene, resulting in high expression levels and improved cell viability compared with electroporation-based methods. In addition, TRAC locus-specific targeted integration was achieved more efficiently through end-joining–based repair pathways than through homology-directed repair following LNP delivery. The resulting engineered CAR-T cells exhibited potent and antigen-specific cytotoxic activity. Together, these results demonstrate that NLS–assisted LNP delivery overcomes a key bottleneck in non-viral gene integration and provides a robust strategy for the generation of functional CAR-T cells.

## Introduction

CAR-T technology has delivered groundbreaking therapeutic effects for refractory hematologic cancers, becoming a game-changer in cancer treatment ^1–3^. Recently, CAR-T technologies have expanded beyond hematologic malignancies, but also solid tumors and autoimmune diseases and its widespread adoption is anticipated ^4,5^. In many ex-vivo CAR-T therapies, CAR transgenes are introduced using lentivirus vectors. However, this approach poses difficulties, such as the high cost of viral production^6^ and an increased risk of secondary tumors due to random transgene integration^7^. Furthermore, gene delivery via electroporation (EP) causes significant damage to cells, leading to issues such as reduced cell yield^8,9^.

Consequently, lipid nanoparticles (LNPs) are an attractive non-viral nucleic acid delivery system that is safer and causes less damage to cells^10^. LNPs can be manufactured cost-effectively and at scale, can encapsulate relatively large cargos, and can facilitate cellular uptake via endocytic pathways. Various applications of LNPs to both ex-vivo and in-vivo CAR-T have begun to be reported. However, most reported LNP-based CAR-T approaches have focused on transient mRNA delivery, reflecting persistent barriers to DNA nuclear entry and stable genomic integration^11^. This reflects several technical barriers. Primary T cells are intrinsically difficult to transfect or transduce, and efficient gene delivery typically requires activation while most circulating T cells remain quiescent. In addition, efficient DNA delivery by LNPs and subsequent genomic integration remain challenging. In general, relatively high integration efficiencies up to 50-60% have been reported for random integration mediated by viral vectors or electroporation, as well as semi-random integration mediated by enzymes such as transposases or large serine integrases^12,13^. However, site-specific integration utilizing homology directed repair (HDR) following genome cleavage by CRISPR/Cas9 is particularly difficult with non-viral methods^14^, and CAR gene integration targeting the TRAC locus is known to be especially inefficient^15^.

We have previously focused on ionizable lipids as key components of LNPs and have developed a proprietary lipid library^16^. Leveraging this expertise, we identified lipids suitable for RNA and DNA delivery into human primary T cells and developed a corresponding LNP formulation. Notably, our newly developed LNP does not require specialized equipment for nucleic-acid encapsulation. Instead, it utilizes a ready-to-use (RtoU) empty LNP technology, allowing users to encapsulate and formulate any nucleic acid into the LNP at any time.

In this study, we aimed to deliver DNA into human primary T cells using our proprietary LNP platform and to achieve efficient integration of a CAR transgene into the host genome.

## Result

### Ready-to-Use LNPs can efficiently deliver mRNA and pDNA to primary T cells and induce transient expression

First, we prepared RtoU LNPs that do not contain nucleic acids (Supplementary Fig. 1A). EGFP mRNA or plasmid DNA (pDNA) was encapsulated in RtoU LNP-R, and the percentage of cells expressing EGFP after transfection into activated human primary T cells, as well as the cell proliferation rate 24 hours after transfection, were evaluated (Fig. 1A). The results showed that for mRNA, both LNP and EP exhibited nearly compatible % EGFP+ cells of approximately 100%, whereas for pDNA, EP showed an 82.6 ± 2.3 % EGFP+ cells compared to 14.6 ± 4.7 % with LNP-R (Fig. 1B and D). Furthermore, regarding proliferation, transfection with LNP-R demonstrated significantly higher rates than EP for both mRNA and pDNA (Fig.1 C and E). There was no significant variation in transfection efficiency or cell proliferation among the three T-cell donors when using LNP. When using LNP-D, which exhibits superior pDNA delivery, the expression rate of the EGFP pDNA was 35.7 % (Supplementary Fig. 1B and C). Additionally, by encapsulating CRISPR Cas9 mRNA and gRNA targeting TRAC into RtoU LNP-R, we were able to achieve gene knockout at a rate of over 90%, which is nearly same level as EP (Fig. 1F). On the other hand, the cell proliferation rate was significantly higher in the LNP-treated group compared to the EP (Fig.1 G). Consequently, the yield of knockout cells was highest in the LNP-treated group (Fig. 1H). Furthermore, in T cells treated with LNP, the percentage of cells positive for γH2AX, an indicator of DNA damage, was as low level as that in untreated cells, indicating that LNP treatment causes significantly less cellular damage compared to EP (Fig. 1I).

**Figure. 1.**
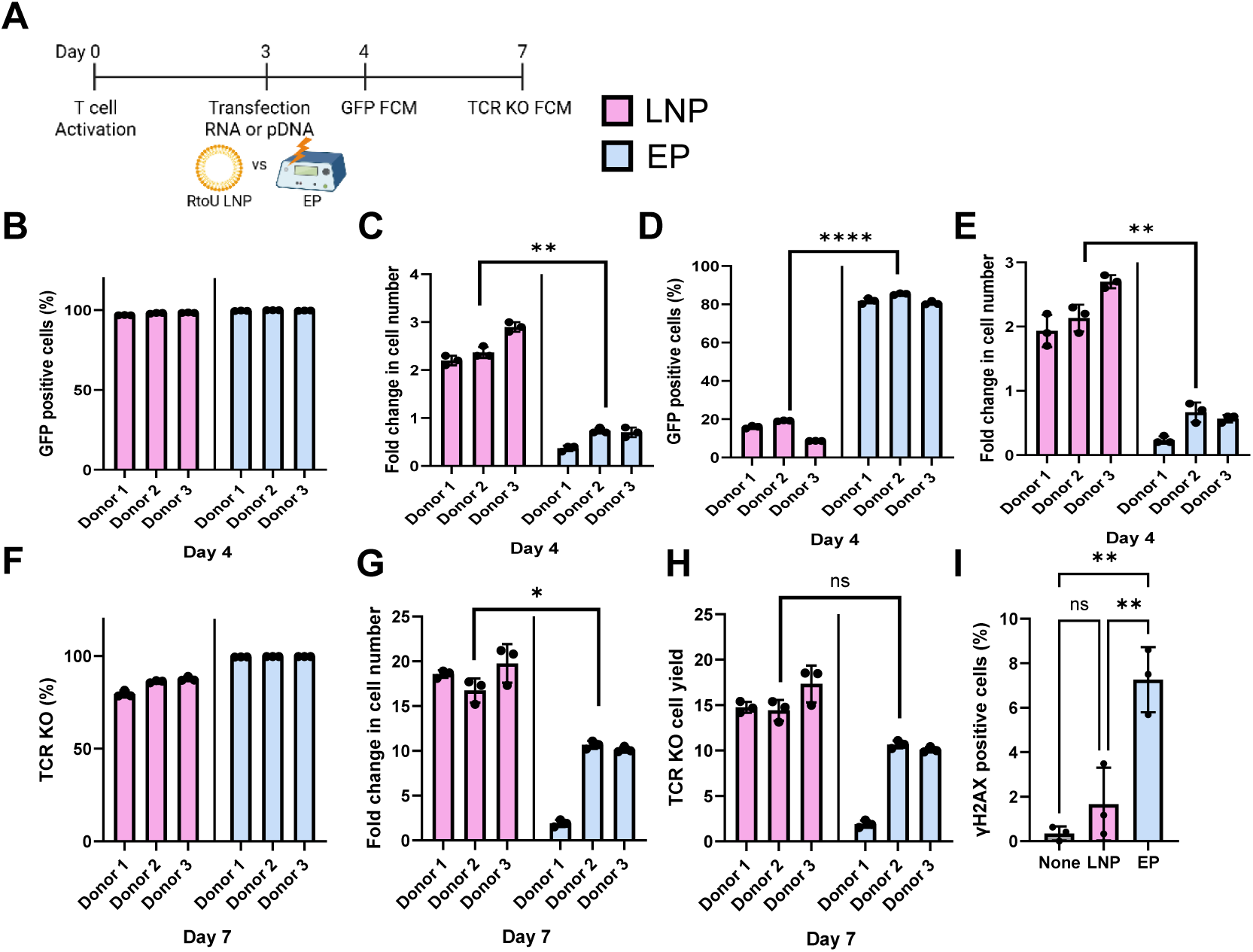
Experimental Schedule for T-Cell Transfection using RtoU LNP or EP (A). FCM represents flow cytometry. EGFP positivity rate (B) and proliferation rate (C) the day after EGFP mRNA transfection into activated human primary T cells using LNP-R (2μg/mL) and electroporation (EP, 6.7μg/mL). EGFP positivity rate (D) and proliferation rate (E) 4 days after EGFP pDNA transfection using LNP-D (2μg/mL) and EP (6.7μg/mL). Mean ± SD of n = 3. Human primary T cells were activated on Day 0, and on Day 3, Cas9 mRNA (LNP; 1.6μg/mL, EP; 5.3μg/mL) and sgRNA (LNP; 0.4ug/mL, EP; 1.3ug/mL) targeting the TRAC locus were transfected using LNP-R or EP. TCR knock-out (KO) efficiency (F), proliferation rate (G), and yield of KO cells (H) on Day 7. Proportion of γH2AX-positive cells at 2 hours post-transfection (I). Mean ± SD of n = 3.

### Lipid Nanoparticles Exhibit Lower Nuclear Translocation of DNA Compared to Electroporation

To compare the intracellular DNA uptake between LNP-D and EP, fluorescein-labeled pDNA (2.7 kbp) was transfected into activated human T cells, and fluorescence was measured at various time points post-transfection. The results showed that the intracellular fluorescein signal increased gradually from 3 to 24 hours after LNP transfection, whereas EP-treated cells exhibited a peak fluorescence at 3 hours followed by a decline over time (Fig. 2A). These findings indicate that while LNP-mediated DNA uptake occurs more slowly, DNA is progressively internalized into cells. Next, we transfected cells with pDNA encoding EGFP (3.5 kbp) to compare protein expression levels. At both 3 and 24 hours post-transfection, EP showed significantly higher EGFP expression compared to LNP-D. Although 25.3 ± 4.8 % EGFP-positive cells were observed in the LNP group at 24 hours, the median fluorescent intensity of EGFP was markedly lower than that seen with EP (Fig. 2B, C, D). Microscopic examination of cells 3 hours after transfection with fluorescein-labeled pDNA revealed distinct localization patterns. LNP-treated cells showed accumulation of pDNA predominantly in the cytoplasmic endosome, whereas EP-treated cells demonstrated nuclear localization of the DNA (Fig. 2E). Consistent with this, droplet digital PCR (ddPCR) analysis of nuclear delivery efficiency at 3 and 24 hours post-transfection confirmed significantly lower nuclear translocation of pDNA following LNP treatment compared to EP (Fig. 2F). Collectively, these results suggest that while LNP-mediated nucleic acid delivery to the cytoplasm is effective, the subsequent translocation of DNA from the cytoplasm into the nucleus is substantially less efficient than that achieved by EP. This difference likely arises because EP physically disrupts the nuclear membrane via electrical pulses, facilitating immediate and direct nuclear delivery of DNA. In contrast, LNPs rely on cellular processes such as membrane fusion, endocytosis, and endosomal escape before DNA can reach the nucleus, and lack an active mechanism for nuclear translocation of nucleic acids.

**Figure. 2.**
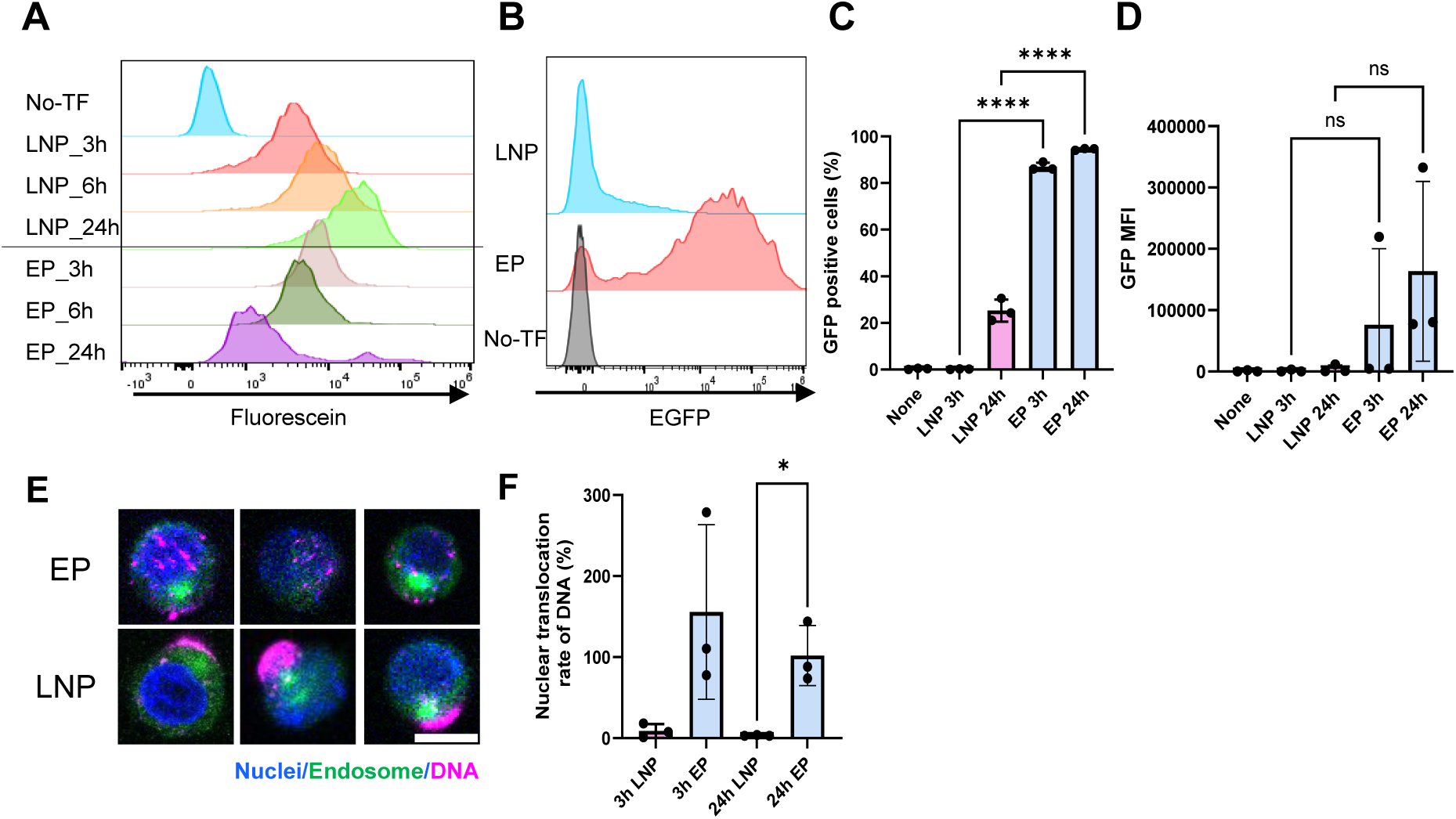
Human primary T cells were activated on Day 0, and on Day 3 fluorescein labeled plasmid DNA was transfected using LNP-D (2μg/mL) or EP (6.7μg/mL). Intracellular fluorescein fluorescence intensity at each time point (A). EGFP expression at 24 hours after EGFP pDNA transfection using LNP-D (2μg/mL) and EP (6.7μg/mL) (B). Percentage of EGFP-positive cells and median fluorescence intensity(D). Mean ± SD of n = 3. T cells 3 hours after transfection with fluorescein-labeled pDNA via LNP-D or EP. The scale bar represents 10um (E). Percentage of pDNA that has translocated into the nucleus 3 and 24 hours after transfection of fluorescein-labeled pDNA via LNP-D or EP (F).

### Nuclear Localization Signal (NLS)-shuttled PiggyBac Improves CAR-DNA Integration Using Lipid Nanoparticles

As demonstrated in the previous section, the nuclear translocation efficiency of DNA delivered via LNP was significantly lower than that achieved by EP. To enhance the genome integration efficiency of CAR pDNA mediated by LNP, we engineered mRNA encoding HyPBase, a hyperactive mutant of the PiggyBac transposase^17^, fused with a strong nuclear localization signal (NLS) derived from SV40^18^. In the transposase system, including HyPBase, requires the inverted terminal repeat (ITR) sequences into the donor DNA, which are recognized and bound by the transposase. We hypothesized that HyPBase translated in the cytoplasm would bind to the ITR sequences on the donor DNA and facilitate nuclear transport of the donor DNA through the appended NLS on HyPBase. As a control, we also prepared HyPBase mRNA lacking the NLS. Both mRNAs, along with a small plasmid containing the CD19 CAR sequence flanked by ITRs as the donor DNA, were encapsulated in LNP-R and delivered into activated primary human T cells (Fig. 3A: experimental overview and timeline). At day 5 post-transfection, CAR integration efficiency with NLS-HyPBase was 4.4 ± 0.3 %, whereas NLS-tagged HyPBase significantly increased CAR integration efficiency to 10 ± 0.3 % (Fig. 3B). Furthermore, reducing the size of the donor pDNA increased the integration efficiency fivefold, reaching a maximum of 76.4 ± 2.0 % at day 14 (Fig. 3C and D, and Supplementary Fig. 2). In comparison, EP resulted in a high CAR integration efficiency (Fig.3D). However, EP caused a marked decrease in cell proliferation relative to LNP delivery (Fig. 3E), ultimately resulting in a higher yield of CAR-T cells generated via LNP compared to EP (Fig. 3F). This is consistent with data shown in Fig. 1I, indicating that EP induces greater cellular and genomic DNA damage than LNP. Finally, CD19 CAR-T cells generated by LNP delivery demonstrated functional cytotoxic activity when co-cultured with CD19-positive B cell lymphoma cells, NALM6, at a stringent E:T ratio (1:10), confirming their antigen-specific cytotoxicity (Fig. 3G and 3H).

**Figure. 3.**
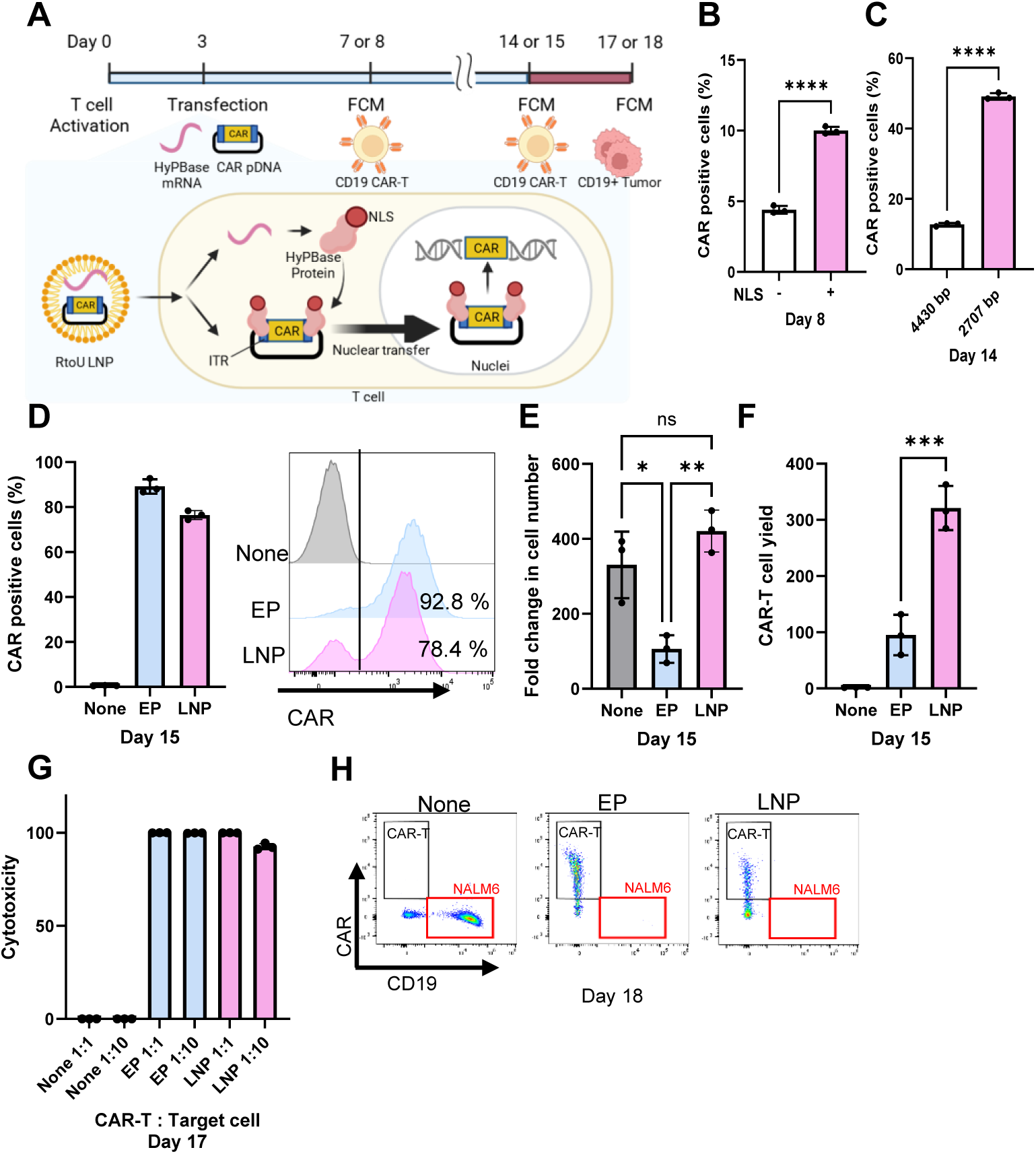
Overview and schematic of the experiment using NLS-shuttled HyPBase transposase. DNA introduced into the cytoplasm by LNP are transported into the nucleus by NLS+ HyPBase, which is translated from mRNA (A). CAR positive rates between NLS- and + HyPBase in human primary T cells 5 days after transfection using the LNP-R. Mean ± SD (n = 3) (B). CAR positive rates between 4,430 bp and 2,707 bp plasmids in human primary T cells 11 days after transfection using the LNP-transposon system. Mean ± SD (n = 3) (C). All subsequent data were obtained using the 2,707 bp plasmid and NLS-fused HyPBase. CAR positive rates between EP and LNP-R in human primary T cells 12 days after transfection. Final nucleic acid concentrations were EP: DNA 5 μg/mL, mRNA 5 μg/mL; LNP: DNA 2 μg/mL, mRNA 1 μg/mL. Mean ± SD (n = 3) (D). Histogram of CAR positive cells under each condition (D, right). Fold change in cell number from transfection between EP and LNP-R in T cells 12 days after transfection. Mean ± SD (n = 3) (E). CAR-T cell yield between the two transfection methods. CAR-T cell yield was calculated by multiplying the CAR positive rate by the fold change in cell number. Mean ± SD (n = 3) (F). Killing assay was performed 12 days after transfection to evaluate the cytotoxicity of CAR-T cells generated by each method. CAR-T cells and target cells (NALM6) were mixed and co-cultured at ratios of 1:1 and 1:10 for 3 days. Cytotoxicity was evaluated by counting the remaining NALM6 cells (G). Representative FACS plots from the killing assay 3 days after co-culture at E:T=1:1 ratio, showing NALM6 cells (CD19) and CAR-T cells (CAR) (H).

### Site-specific CAR Knock-in Efficiency Using CRISPR/Cas9 Is Enhanced by HITI/NHEJ and PITCh /MMEJ Methods with NLS Shuttle System Compared to HDR

Random integration of DNA via viral vectors or semi-random integration mediated by transposases carries the risk of insertion near oncogenes or cell cycle-related genes, potentially leading to secondary tumorigenesis^7^. While site-specific integration is increasingly demanded for safety, the insertion efficiency using the CRISPR/Cas9 system remains low^15^. To address this, we attempted TRAC locus-specific knock-in of a CAR gene using CRISPR/Cas9 combined with LNP-R and LNP-D.

Among various DNA repair pathways employed for DNA insertion via CRISPR/Cas9, we hypothesized that NHEJ-based HITI ^19,20^ and MMEJ^21^-based Precise Integration into Target Chromosome (PITCh)^22,23^ donors bearing Cas9 target sequences (CTS; i.e., the sgRNA recognition site) would associate with NLS-tagged Cas9 in the cytoplasm and thereby enhance donor nuclear entry, yielding higher knock-in efficiency than HDR (Fig. 4A). We prepared NLS-modified Cas9 mRNA along with donor pDNA encoding a CD19-CAR sequence designed for HDR, HITI, or PITCh (Fig. 4A and Supplementary Fig.3A). The NLS-modified Cas9 mRNA was encapsulated into LNP-R and donor pDNAs were encapsulated into LNP-D. They were introduced into activated primary human T cells (Fig. 4A). At day 14, knock-in efficiency by HDR was very low (1.0 ± 0.3 %), whereas HITI and MMEJ achieved substantially higher knock-in efficiencies of 5.7 ± 0.2 % and 19.0 ± 4.1 %, respectively (Fig. 4B and Supplementary Fig. 3B). On the other hand, HITI (16.7 ± 1.6 %) resulted in the lowest CAR knock-in efficiency mediated by EP, while HDR (35.1 ± 1.6 %) ranked second, and MMEJ (55.0 ± 6.0 %) showed the highest efficiency (Fig. 4C). All CD19 CAR-positive cells were negative for TCRαβ expression, supporting site-specific integration at the TRAC locus targeted by the sgRNA (Fig.4F). Regardless of the gene editing method used (HDR, HITI, or PITCh), LNP showed superior post-treatment proliferation compared to EP (Fig.4D and E) and CAR-T cell yield following the PITCh method was comparable between LNP and EP (Fig.4G).

**Figure. 4.**
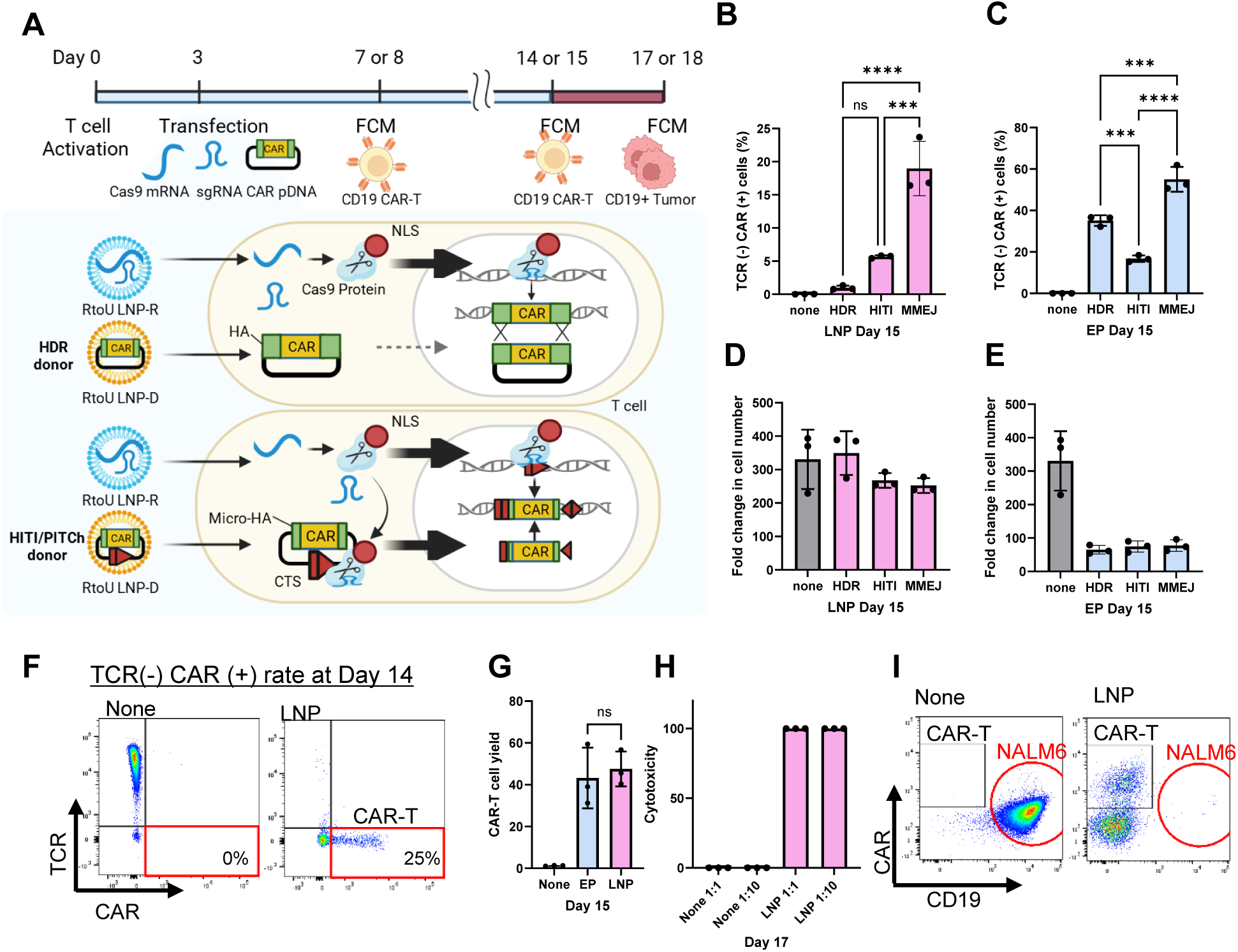
Experimental timeline and CAR gene knock-in (KI) using CRISPR/Cas9. While donor pDNA has difficulty translocating into the nucleus in HDR, HITI and PITCh donor pDNA contain a Cas9 target sequence (CTS). Nuclear translocation of the donor pDNA is induced by binding NLS-Cas9 to donor pDNA. HA represents homology arm (A). TCR-CAR+ rates using indicated LNP-CRISPR KI in human primary T cells 11 days after transfection. Final concentrations were DNA: 2 μg/mL and mRNA: 4 μg/mL. Data represent mean ± SD (n = 3) (B). TCR-CAR+ cell rates using EP in human primary T cells 12 days after transfection. Final concentrations were DNA: 5 μg/mL and mRNA: 5 μg/mL. Mean ± SD (n = 3) (C). Comparison of CAR-T cell yield between the two transfection methods using PITCh vector by MMEJ pathway. CAR-T cell yield was calculated by multiplying the TCR-CAR+ rates by the fold change in cell number. Mean ± SD (n = 3) (D). Representative FACS plot of cells transfected via the LNP-CRISPR system (MMEJ), showing the TCR-CAR+ population (E). Fold change in cell number 12 days after transfection (F: LNP, G: EP). Killing assay was performed 12 days after transfection to evaluate the cytotoxicity of CAR-T cells generated by the LNP-CRISPR system using PITCh vector. CAR-T cells and target cells (NALM6) were mixed and co-cultured at ratios of 1:1 and 1:10 for 3 days. Cytotoxicity was evaluated by counting the remaining NALM6 cells using flowcytometry (FCM). Mean ± SD (n = 3) (H). Representative FCM plots from the killing assay 3 days after co-culture, showing NALM6 cells (CD19) and CAR-T cells (CAR-T) (I).

Furthermore, CD19 CAR-T cells generated using PITCh methods demonstrated cytotoxic activity against CD19-positive B cell lymphoma NALM6 cells (Fig. 4H and I). We also observed that separating the delivery of NLS-Cas9 mRNA and sgRNA in LNP-R and donor pDNA in LNP-D showed higher knock-in efficiency compared to co-encapsulation in LNP-R (Supplementary Fig. 3C). This is presumably because co-encapsulation in the RtoU LNPs leads to interference between Cas9 mRNA and donor DNA.

Additionally, converting the donor DNA for HDR from double-stranded small plasmid DNA into circular-single-stranded (css) DNA resulted in an 5-fold increase (cds-HDR 1.0 ± 0.3 %, css-HDR 5.2 ± 0.5 %) in knock-in efficiency, although the efficiencies did not reach those of HITI or PITCh (Supplementary Fig. 3D). Treatment with the DNA-PK inhibitor AZD7648 enhanced HDR-mediated knock-in efficiency up to 4-fold, but this was accompanied by a dose-dependent decrease in cell proliferation (Supplementary Fig. 3E).

Overall, while HITI and PITCh exploit NHEJ or MMEJ pathways that are active in non-dividing cells, our results suggest that, under LNP-mediated delivery conditions, donor DNAs containing CTS used in HITI and PITCh appear to benefit from an NLS shuttle–mediated enhancement of nuclear entry relative to HDR donors lacking CTS.

## Discussion

In this study, we first demonstrated that RtoU LNPs can efficiently deliver RNA into primary human T cells with high efficiency and minimal cellular damage. Subsequently, we evaluated pDNA delivery efficiencies, achieving approximately 30% with LNP-R and up to 70% with LNP-D. When comparing DNA uptake between LNP and EP, although there was a temporal lag in intracellular DNA uptake, both methods successfully introduced DNA into the cells. However, protein expression following LNP-mediated delivery was markedly lower, which we believe results from restricted nuclear translocation of the introduced DNA.

One reason why DNA-delivering LNPs have not been widely developed is that the process requires two steps, escape from endosomes and nuclear translocation of the DNA. We identified ionizable lipids and LNPs with high endosome-escape capacity, achieving transient expression of pDNA up to 40% (LNP-D). However, since DNA nuclear translocation is a phenomenon independent of LNP or constituent lipids, it was necessary to modify the nucleic acid component. To address this, we focused on introducing NLS, which are short peptide sequences that actively mediate transport from the cytoplasm to the nucleus by binding to cytoplasmic receptors such as importin β ^24^. Direct conjugation of NLS peptides to DNA or encapsulation within RtoU LNPs suggested technically challenging. Therefore, we devised a shuttle system wherein NLS sequences were fused to mRNAs encoding enzymes such as transposase or Cas9, alongside donor DNA containing the corresponding enzyme recognition sites. This enabled the NLS-tagged enzyme proteins to bind donor DNA within the cytoplasm and facilitate active nuclear translocation of the complex.

In the transposase HyPBase system, which inherently exhibits relatively high CAR transgene integration efficiencies (10–30%), the introduction of the NLS shuttle system combined with reduction of donor DNA size considerably enhanced integration efficiency, reaching approximately 80%. It has been reported that when using EP to insert PiggyBac mRNA and donor CAR pDNA into T cells, the copy number increases in proportion to the amount of nucleic acid^25^. With LNP, the nuclear pDNA abundance may be lower than with EP, consistent with our nuclear delivery measurements (Fig. 2F). Furthermore, since the transposon method preferentially inserts into TTAA sequences in the genome, the risk of insertional mutagenesis is considered to be lower than that associated with retroviral vectors.

Similarly, in CRISPR knock-in system, the NLS shuttle markedly improved knock-in rates. For instance, HDR donors with long homology arms lacking a CTS achieved knock-in efficiencies below 1.5 %, whereas donor DNAs incorporating CTS for NHEJ and MMEJ pathways reached efficiencies of 5.7 ± 0.2 % and up to 25 %, respectively. In addition to the NLS shuttle, the size of the donor pDNA is another key factor affecting insertion efficiency. While insertion efficiency increases as the size decreases, in practice, 3 kbp is the critical threshold. Importantly, CAR-T cells generated using virus-free LNP-mediated methods demonstrated robust tumor cell cytotoxicity, confirming their functional competence.

Taken together, our results represent a significant advance in the efficiency of site-specific gene integration in T cells using LNP delivery, improving efficiencies up to 25 %, a notable achievement given the traditionally low performance of LNP-based gene targeting. Targeted CAR knock-in at the TRAC locus remains a well-established approach to enhance CAR-T cell activity^26^. Recently, hybrid methods combining PiggyBac transposons with CRISPR-Cas9 have emerged for site-specific integration; however, the disruption of PiggyBac’s transposition activity via genetic mutation appears to limit efficiency^27^.

In summary, our NLS shuttle strategy leveraging the NHEJ or MMEJ pathway offers a promising platform for enhancing gene knock-in efficiencies with LNPs. This approach holds substantial potential for diverse applications in ex-vivo and in-vivo gene and cell therapies, including the development of virus-free CAR-T cells.

## Materials and Methods

### RtoU LNP production

The ionized lipids and other reagents used, as well as the synthesis method, were prepared as described in the patent^28^. An aqueous solution at pH = 4.0 was rapidly mixed with an ethanol solution of lipids then dialyzed and concentrated, thereby obtaining lipid particles devoid of nucleic acids (RtoU LNPs). LNPs used in this study contained a proprietary mixture of ionizable lipid, phospholipid, cholesterol, and PEG-lipid. Based on differences in the ionizable lipids, we developed LNP-R, which has high mRNA delivery efficiency, and LNP-D, which has high DNA delivery efficiency, and used them in subsequent experiments.

### Nucleic acids

EGFP mRNA, CleanCap Cas9 mRNA (5moU) (TriLink, L-7206), Label IT Plasmid Delivery Control, Fluorescein (Takara Bio, V7906) were purchased. sgRNA targeting the human T-cell receptor alpha constant (TRAC) gene (target sequence: A*G*A*GUCUCUCAGCUGGUAACA, ThermoFisher, A35514) was synthesized. EGFP pDNA (pMAXGFP, Lonza) was included with P3 Primary Cell 4D-Nucleofecto X Kit L (Lonza, V4XP-3012). The Hyperreactive PiggyBac (HyPBase) sequences and NLS (+) HyPBase sequences were synthesized as mRNA by Genscript, based on the reported data^17^. NLS (+) HyPBase sequences were designed by adding the SV40 NLS on the both termini. The donor template for the small plasmid CD19CAR insertion refers to the report ^15^, small GenCircle DNA constructs were synthesized by Genscript.

For the sequence of CD19 CAR and the homology arms used to insert the CAR transgene into the TRAC locus for HDR, refer to the report^15^, small GenCircle DNA constructs were synthesized by Genscript.

For circular single-stranded DNA (cssDNA), the donor plasmid for the HDR pathway (double-stranded DNA, dsDNA) was reacted with the restriction enzyme Nb.Bpu10I (ThermoFisher Scientific, ER1681) at 37°C for 1 hour to introduce nicks into the plasmid. After purifying this using the QiAcquick PCR Purification Kit (Qiagen), Exonuclease III (Thermo Fisher Scientific) was added and the mixture was incubated at 30°C for 10 minutes to cleave the nick-containing single-stranded DNA, thereby preparing circular single-stranded DNA (cssDNA). Furthermore, the resulting circular single-stranded DNA was purified again using the QiAcquick PCR Purification Kit (Qiagen). The donor pDNA sequences for HITI^19^ and MMEJ^21^ were modified based on previous papers and synthesized as GenCircle DNA by Genscript. The relevant nucleic acid sequences are listed in Supplemental Table 1.

### Preparation of mixture RtoU LNP and nucleic acids

Freeze-dried nucleic acids were reconstituted in UltraPure DNase/RNase-Free Distilled Water (Invitrogen) to prepare 1-2 mg/mL solution. Empty LNP stored at −80°C was thawed at room temperature. In this experiment, 0.8 mg/mL of nucleic acid, LNP, and neutralizing solution were mixed in a mass ratio of 1:2:5. After mix of nucleic acids and LNP solution at room temperature for 5 minutes, an appropriate volume of neutralizing solution (20 mmol/L Tris buffer (pH 8.4) containing 8% sucrose) was added and combined by pipetting to prepare nucleic acid-encapsulated LNPs. The neutralized nucleic acid-LNP mixture was added to the cell culture medium to achieve the indicated final nucleic acid concentration.

### Primary cell culture

Human primary T cells (Human PB Pan-T, Cryo; STEMCELL Technologies, ST-70024), derived from healthy adult donors with informed consent, were purchased from STEMCELL Technologies. This study was approved by the Institutional Review Board of BioScience and Engineering Laboratories, FUJIFILM Corporation (approval no. 2022-005). Frozen human primary T cells were thawed by incubating them in a water bath at 37°C for several minutes. The thawed T cells were resuspended in PRIME-XV T cell CDM　(FUJIFILM Biosciences), washed by centrifugation, and then resuspended in T-cell culture medium (PRIME-XV T cell CDM (FUJIFILM Biosciences) containing 10 ng/ mL of hIL-2 (Roche), 5 ng/mL of hIL-7, and hIL-15 (Miltenty)). T cells were prepared at a concentration of 1.0 × 10^6^ cells/mL using T-cell culture medium and resuspended in DynaBeads Human T-Activator CD3/CD28 (Thermo Fisher, DB11131) were added at a 1:1 bead to T cell ratio. The cells were seeded into 24-well plates for cell culture and activated by culturing them in a 37°C, 5% CO_2_ incubator for 3 days. On the third day of activation, the DynaBeads were removed from the T cell culture medium. The T cells pretreated in this manner were used as activated T cells.

### In-vitro transfection

#### LNP

The activated T cells were adjusted to a concentration of 1.0 × 10^6^ cells/mL in T-cell culture medium containing recombinant human apolipoprotein E3 (ApoE3) (Fujifilm Wako, 010-20261) at a final concentration of 1 μg/mL, as prepared in advance, and seeded into a 96-well plate. Using LNPs encapsulating the mRNA or donor DNA prepared as described above, we added mRNA and/or DNA to achieve indicated final concentration per 1.0 × 10^5^ cells, and cultured the cells in a 37°C, 5% CO_2_ incubator. LNP uptake was measured using LNPs loaded with pDNA conjugated to Fluorescein (Takara bio, V7906).

#### EP

Activated primary human T cells were resuspended, pooled, and washed twice in sterile PBS at 100*g* for 10 min, RT. Afterward, they were resuspended in 20 μL/4×10^5^ cells ice-cold electroporation buffer (P3 [Lonza] as indicated). The exposure time to the electroporation buffers was kept as short as possible. For electroporation of 4 × 10^5^ cells, 20 μL of resuspended cells were transferred to nucleic acids and mixed thoroughly. Afterward, the T cell/nucleic acids mixture was transferred onto a 16-well electroporation strip (20 μL = 4 × 10^5^ cells per well, Lonza). Electroporation was performed on a 4D-Nucleofector Device (Lonza) using the program FI-115. Transfected cells were transferred to 96-well plates containing 200 μL pre-warmed T cell medium per well.

### Cell lines

Nalm6 (CRL-3273) cell lines were procured from ATCC. Nalm6 were cultured in RPMI1640 medium (Thermo Fisher, A1049101) supplemented with 10% fetal bovine serum (FBS, Gobco, 0437-028).

### Flow cytometry

Transfected T cells were harvested, stained with Fixable Viability Stain 780 (BD Biosciences, 565388), and blocked with Human BD Fc Block (BD Biosciences, 564219). CD19 CAR expression was detected using FITC-labeled Anti-FMC63 scFv antibody (AcroBioSystems, FM3-FY45). Additionally, in the CRISPR knock-in assay, TCR expression was evaluated by staining cells with the Anti-Human TCRαβ antibody, PE-labeled (BD Biosciences, 564728). In the in vitro killing assay, after measuring the CAR-positive rate using the above staining method, the required number of CAR-T cells was co-cultured with NALM6 cells. After 3 days, the cells were stained with FVS780, FcBlock, and Anti-FMC63 scFv antibody FITC-labeled (AcroBiosystems, FM3-FY45), and PE Mouse Anti-Human CD19 (BD Biosciences, 555413), and detected by flow cytometry. Flow cytometry analysis was performed using a Attune CytPix flow cytometer (ThermoFisher). Data analysis was performed using FlowJo software (BD Biosciences).

### ddPCR

Cells were suspended with PBS supplemented with sodium heparin (Sigma Aldrich, H3393) and incubated for 30min at 37℃. The remaining nuclear pellets were resuspended in digestion buffer containing Benzonase nuclease (e.g., 25–50 U/mL) and 2 mM Magnesium chloride and incubated for 45min at 37℃. Next, we isolated nuclear fractions using the EPIXTRACT Nuclear Protein Isolation Kit (Enzo, ENZ-45016) according to the instructions. Genomic DNA was purified from the nuclear pellet using the QIAamp DNA Mini Kit (QIAGEN, 51306) following the instructions. Absolute quantification of nuclear pDNA was performed using the ddPCR Supermix for Probes (Bio-Rad Laboratories) and the QX200 Droplet Digital PCR system. The reaction mixture (20 µL) contained 1 x ddPCR Supermix for Probes, 900 nM of each primer, 250 nM of the probe, and 10 ng of template DNA. Droplets were generated using a QX200 Droplet Generator, and PCR amplification was performed with the following conditions: 95℃ for 10 min, followed by 40 cycles of 94℃ for 30 s and 55℃ for 1 min, and a final step at 98℃ for 10 min. After PCR, droplets were analyzed using the QX200 Droplet Reader and QuantaSoft software.

### Immunofluorescence

Fluorescein-labeled pDNA was transfected into T cells via LNP or EP, and the cells were harvested at indicated time points post-treatment. The cells were fixed by incubating them in eBioscienc IC Fixation Buffer (Thermo Fisher, 00-8222-49) at 4 ℃ for 30 minutes. The cells were washed with PBS containing 2% FBS, then stained by adding RAB11A Recombinant Rabbit Monoclonal Antibody (3H18L5), Alexa Fluor 488 (Thermo Fisher Scientific, MA7-00184-A488, 1:100) and incubating at 4℃ for 1 hour. Subsequently, the washed cells were placed on a slide, mounted with Fluoromount-G Mounting Medium, with DAPI (Invitrogen, 00-4959-52), covered with a coverslip, and observed and imaged using a confocal microscope using ECLIPSE Ti (Nikon).

### Functional T cell assays

A flowcytometry-based killing assay was performed to evaluate the functionality of LNP engineered CAR T cells against CD19+ target cells. Prior to the assay, CAR expression was measured by flowcytometry and only CAR+ T cells were counted as effectors. On day 14 post-transfection, 10,000 CAR T cells were co-cultured with target cells Nalm6 at varying E:T ratios (1:1 and 1:10) for 72 hours. After co-culture, the cells were stained with Fixable Viability Stain 780 (BD,565388), FITC-labeled monoclonal anti-FMC63 antibody (Acrobiosystems, FM3-FY45), and PE Mouse Anti-Human CD19 (BD, 555413), and the number of live cells in each well was measured by flow cytometry (Attune CytPix, Thermo Fisher Scientific). Cytotoxicity was calculated using the following formula.

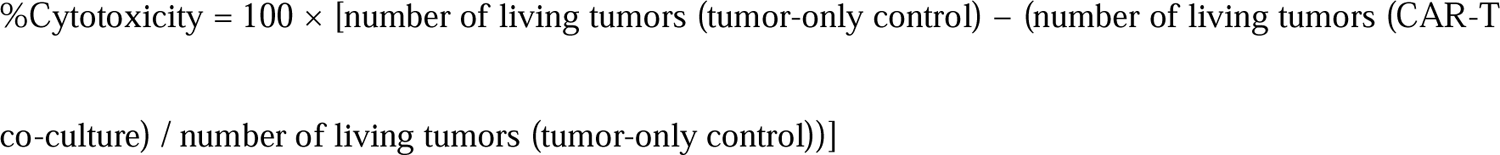

### Statistical analysis

All statistical analyses and graph generation were performed using Prism (GraphPad Software Inc., San Diego, CA, USA). Results are expressed as mean ± standard deviation (mean ± SD). Welch’s t-test was used for comparisons between two groups with n≧3. A standard one-way analysis of variance (ANOVA) was performed. Where significant differences were detected, Tukey’s multiple comparison test was conducted. A p-value less than 0.05 was considered statistically significant (* p<0.05, ** p<0.01 and ***p<0.001, ****p<0.0001).

### Illustration

The illustration was created using BioRender.

## Acknowledgements

The authors thank Sho Toyonaga, Kaori Takeda, Keiichi Onodera, Toshifumi Kimura, Yukiko Ishii and Takeshi Yamamoto for technical assistance and helpful discussions.

## Author Contributions

N.S.-Y. and K.I. and S.U. and Y.M. conceived the study. H.F. and S.H. designed the lipids and developed the RtoU LNP concept. K.S. and K.Y. and K.N. made the RtoU LNPs. K.I. and M.M. and S.U. and K.N. performed experiments using human primary T cells and analyzed data. N.S.-Y. and K.I. and M.M. wrote the manuscript with input from all authors.

## Declaration of generative AI and AI-assisted technologies in the manuscript preparation process

During the preparation of this work, the authors used Microsoft 365 Copilot in order to improve the clarity, grammar, and scientific writing style of the manuscript. After using this tool, the authors reviewed and edited the content as needed and takes full responsibility for the content of the published article.

---

**Supplementary Fig. 1.**
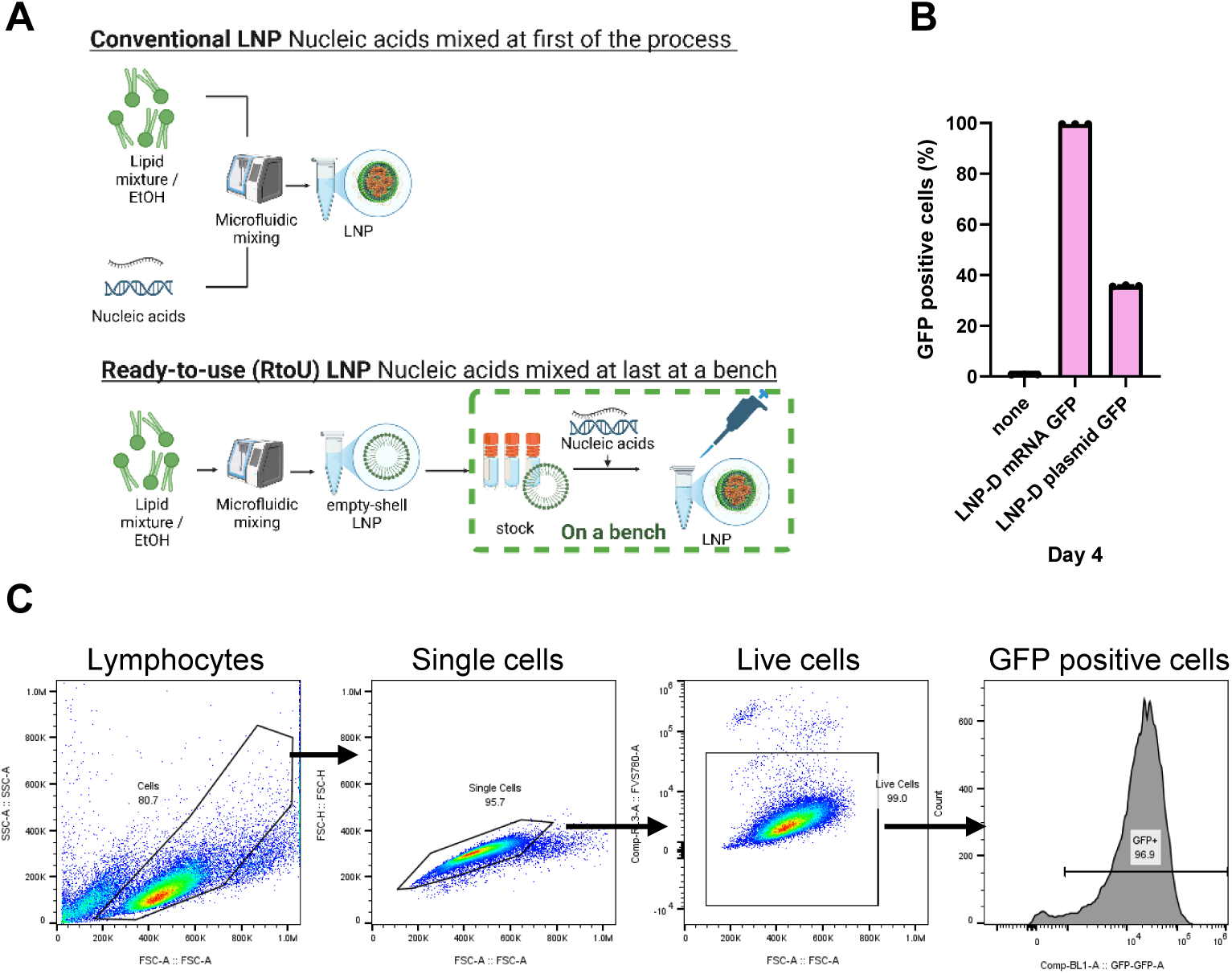
Conventional LNP preparation method and Ready-to-Use LNP (A). GFP positive rates of T cells transfected with LNP-D using mRNA GFP and plasmid GFP (B). Calculation of the percentage of GFP-positive cells following sequential gating of lymphocytes, singlets, and live cells by flow cytometry analysis (C).

**Supplementary Fig. 2.**
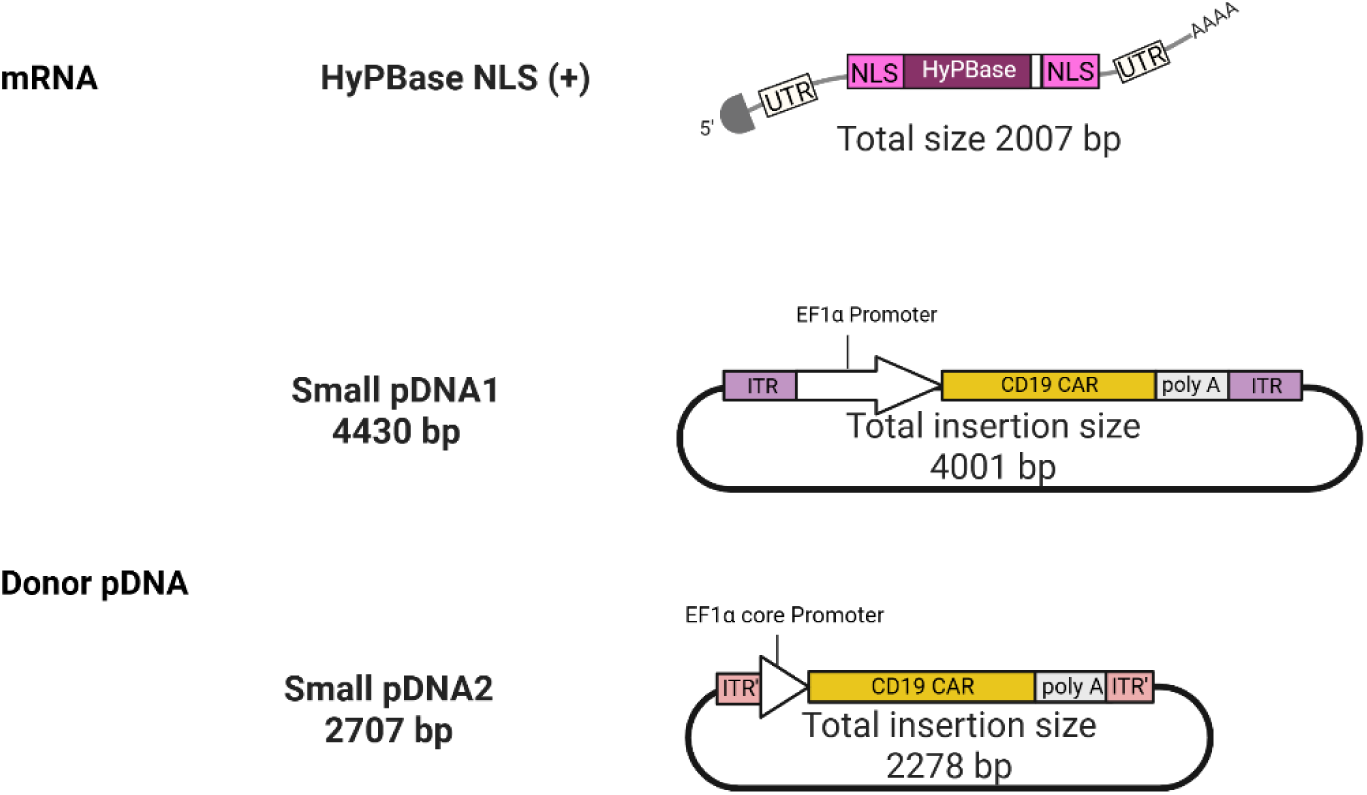
NLS-tagged HyPBase mRNA and donor vector construct for CAR-DNA integration by transposon system. Small pDNA1 contains ITR (Inverted terminal repeat), EF1α promoter, CD19 CAR, and polyA. In small pDNA2, ITR sequence was changed more smaller and EF1α core promoter instead of normal promoter.

**Supplementary Fig. 3.**
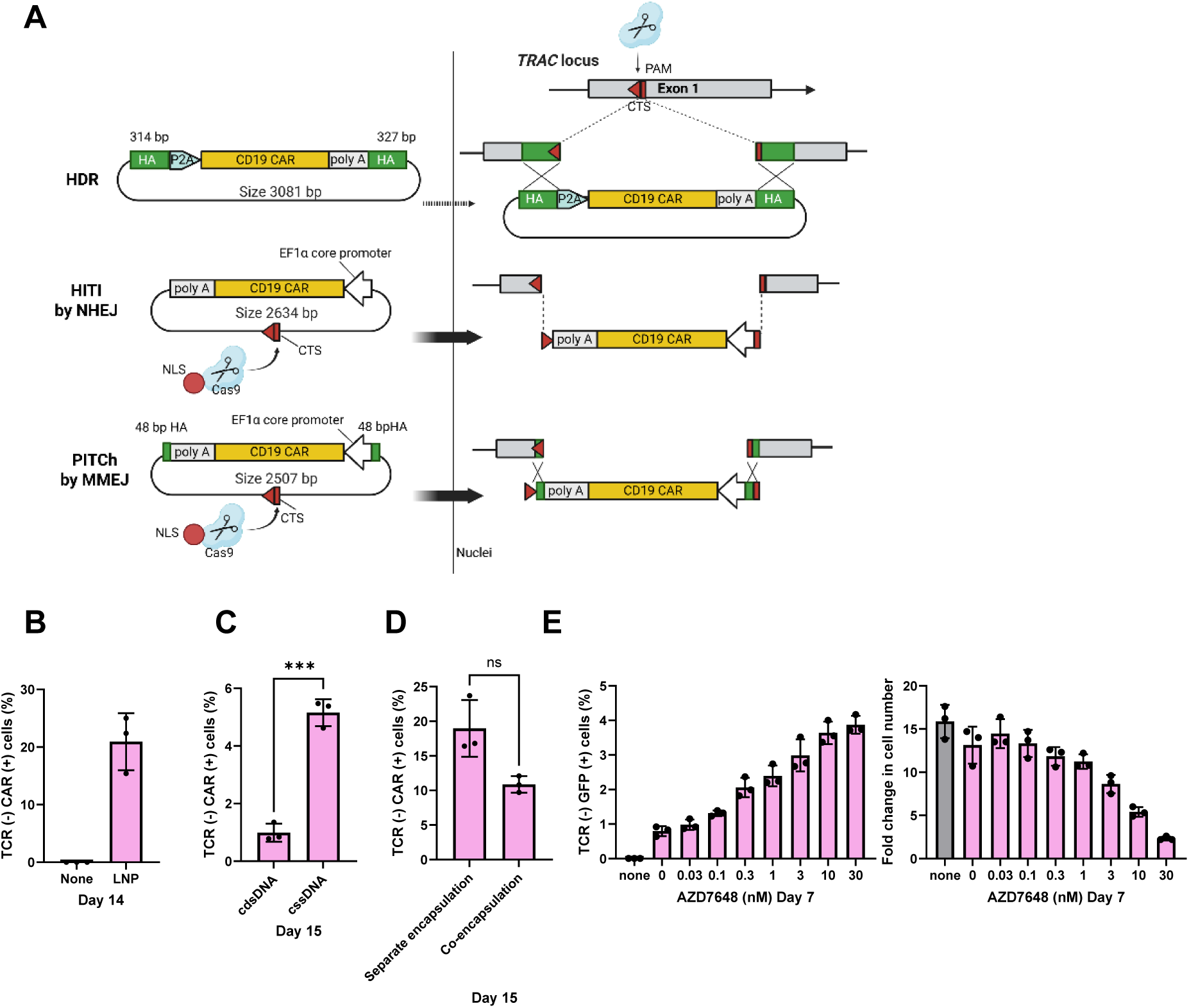
CAR-DNA knock-in (KI) strategy using CRISPR/Cas9. Vector constructs for HDR, HITI/NHEJ, and PITCh/MMEJ, and methods for knock-in into the TRAC locus of the host genome. HA represents the homology arm, and CTS represents the Cas9 (gRNA) target sequence (A). Maximum KI efficiency of the LNP-CRISPR/Cas9 system using PITCh vector by MMEJ. The graph shows TCR-CAR+ cell rates 11 days after transfection (B). TCR-CAR+ cell rates using the LNP-CRISPR/Cas9 system between separate encapsulation of mRNA and DNA in two different LNPs and co-encapsulation of mRNA and DNA in a single LNP (C). TCR-CAR+ cell rates using the LNP-CRISPR/Cas9 system between circular double-stranded DNA (cdsDNA) and circular single-stranded DNA (cssDNA) 12 days after transfection (D). Concentration evaluation of the HDR enhancer AZD7648 using the LNP-CRISPR/Cas9 system (HDR) 4 days after transfection, showing TCR-GFP+ rates (D, right) and fold expansion (D, left).

