## Supplemental Table 1 for "Active Nuclear Shuttling Enables Efficient Virus-Free CAR Gene Integration Using Ready-to-Use Lipid Nanoparticles"

| sequence name | size (bp) | sequence |
|---|---|---|
| HyPB CAR small plasmid 4430 bp | 4430 | [illegible] |
| HyPB CAR small plasmid 2707 bp | 2707 | [illegible] |
| NLS (+) HyPBase | 1842 | [illegible] |
| CRISPR-HDR CAR | 3081 | [illegible] |
| CRISPR-HDR GFP | 2076 | [illegible] |
| CRISPR-HITI CAR | 2634 | [illegible] |
| CRISPR-MMEJ CAR | 2507 | [illegible] |
